# Enhanced beta power emerges from simulated parkinsonian primary motor cortex

**DOI:** 10.1101/2024.05.23.595566

**Authors:** Donald W Doherty, Liqiang Chen, Yoland Smith, Thomas Wichmann, Hong-Yuan Chu, William W Lytton

**Affiliations:** Department of Physiology & Pharmacology, SUNY Downstate Medical Center, Brooklyn, NY 11203, USA; Aligning Science Across Parkinson’s (ASAP) Collaborative Research Network, Chevy Chase, MD, 20815; Department of Pharmacology and Physiology, Georgetown University Medical Center, Washington D.C., USA; Emory National Primate Research Center, Department of Neurology, Udall Center of Excellence for Parkinson’s Disease Research, Emory University, School of Medicine, Atlanta GA 30329 USA; Kings County Hospital, Brooklyn, NY 11203, USA

## Abstract

Primary motor cortex (M1) layer 5B pyramidal tract (PT5B) neurons develop intrinsic pathology in rodent and primate Parkinson’s disease (PD) models. We used computer simulation to predict how decreased PT5B neuron excitability, identified with current injection in vitro, would change activity patterns of the M1 network. Using NEURON/NetPyNE, we implemented computer simulations of PT5B neurons based on control and 6-OHDA-treated mouse slice data. Parkinsonian PT5B neurons, in an otherwise unmodified simulated M1 network, produced major changes in LFP oscillatory power: an order of magnitude increase in beta band power around 15 Hz in the rest state. This demonstrated that relatively small changes in PT5B neuron excitability would alter oscillatory patterns of activity throughout the M1 circuit, increasing beta band power, a signature of PD pathophysiology. Dysfunction in PT5B neurons, the final-common-pathway to brainstem and spinal cord, provides a new target to treat PD motor symptoms.

## Introduction

Motor signs and symptoms of Parkinson’s disease (PD) are causally linked to the degeneration of dopaminergic neurons in the substantia nigra pars compacta (SNc) and their striatal projections.^1^ This pathologic change results in profound changes within the basal ganglia-thalamocortical network of connections. The traditional view, the Delong-Albin model, identified SNc degeneration as the only major pathology; in this view, the nigrostriatal projection loss created an imbalance in the ‘direct’ vs. ‘indirect’ pathways of the basal ganglia (BG) so that the overall-negative indirect route dominated, giving reduced cortical activity.^2–4^ Other studies indicated the importance of oscillations, showing that DA loss promoted the emergence of movement-suppressing beta band activity in the cortico-basal ganglia-thalamocortical network and the reduction of gamma band oscillations.^2,5,6^ At the single cell level, these changes are detectable as synchronized oscillatory burst patterns of neighboring neurons. Oscillatory changes consistently correlate with the severity of PD motor deficits, and are now seen as being more tightly linked to the emergence of parkinsonism than are changes in cortical firing rates predicted by Delong-Albin.

As an additional corrective to the traditional model, it is now apparent that PD pathology does not remain confined to the SNc. At the molecular and cellular levels, loss of SNc DA neurons triggers additional adaptive changes in neurons of other motor areas and in their synaptic connections. For example, the number of corticostriatal and corticosubthalamic terminals decreases, and the functional connectivity between the globus pallidus and the STN increases after the loss of SNc DA neurons.^7–10^ Functionally, DA depletion alters the intrinsic excitability of neurons within the basal ganglia and impairs the synaptic plasticity at the cortico-striatal and cortico-STN inputs.^10–12^ These adaptive changes may play an important role in the development of the abnormally synchronized pattern of activity within the basal ganglia.^5,6,13,14^

The primary motor cortex (M1) is the most prominent source of corticospinal projecting neurons, and is likely to have a disproportionate role in translating pathological neural activity into motor deficits in PD.^15^ Prior *in vivo* studies of MPTP-treated parkinsonian macaques reported changes in the firing rates and patterns of pyramidal neurons in M1, particularly affecting the layer 5B (L5B) pyramidal tract (PT5B) corticospinal neurons, which likely relates to impaired movement encoding under parkinsonian conditions.^16–18^ Our recent work confirmed and extended our knowledge of the structural and functional adaptations of M1 pyramidal neurons in parkinsonian monkeys and rodents, showing selectively reduced intrinsic excitability of the PT5B neurons in parkinsonian mice,^19^ and anatomical and morphological changes in these cells in MPTP-treated parkinsonian monkeys.^20^ Together, these data suggest that, following the loss of DA, M1 becomes another site of dysfunction, where cell-subtype-specific cellular and synaptic adaptations give rise to maladaptations.

Nonlinear dynamical theory of computational neuroscience shows that dynamical dysfunctions will emerge as a consequence of localized changes. We hypothesized that the decreased cellular excitability of PT5B neurons would disrupt firing and oscillatory patterns in the M1 network. In the present work, we tested this hypothesis using computer simulations with our previously validated computer model of mouse M1.^21^ We found that decreased intrinsic excitability of PT5B neurons paradoxically increased their firing activity in the network environment. Our simulations also showed reduced variability in PT5B neuronal firing, and a significant increase in the beta-band power at the population level, correlating with beta-band changes found in PD.

## Results

Biophysically-realistic, empirically-validated simulations utilized a mouse M1 network model composed of more than 10,000 neurons comprising PT5B, IT, PV, SOM neurons that were distributed across 6 cortical layers in a 300 μm diameter cylindrical volume (Fig 1).^21,28^ Recent comparison of model results with in vivo mouse M1 recordings and behavior validated the data driven M1 model’s cell-type specific responses during resting (quiet wakefulness) and activated (movement) behaviors.^21^ In addition, local field potential (LFP) results were validated through the analyses of their sources.^21^ In the present study, over 100,000 simulations were run in developing and exploring these simulations; one second of simulation time took about 2 hours to compute on a 5.16 peak petaflops 64-node supercomputer. The simulations were done under resting and activated states, as defined above.^21^

**Figure 1.**
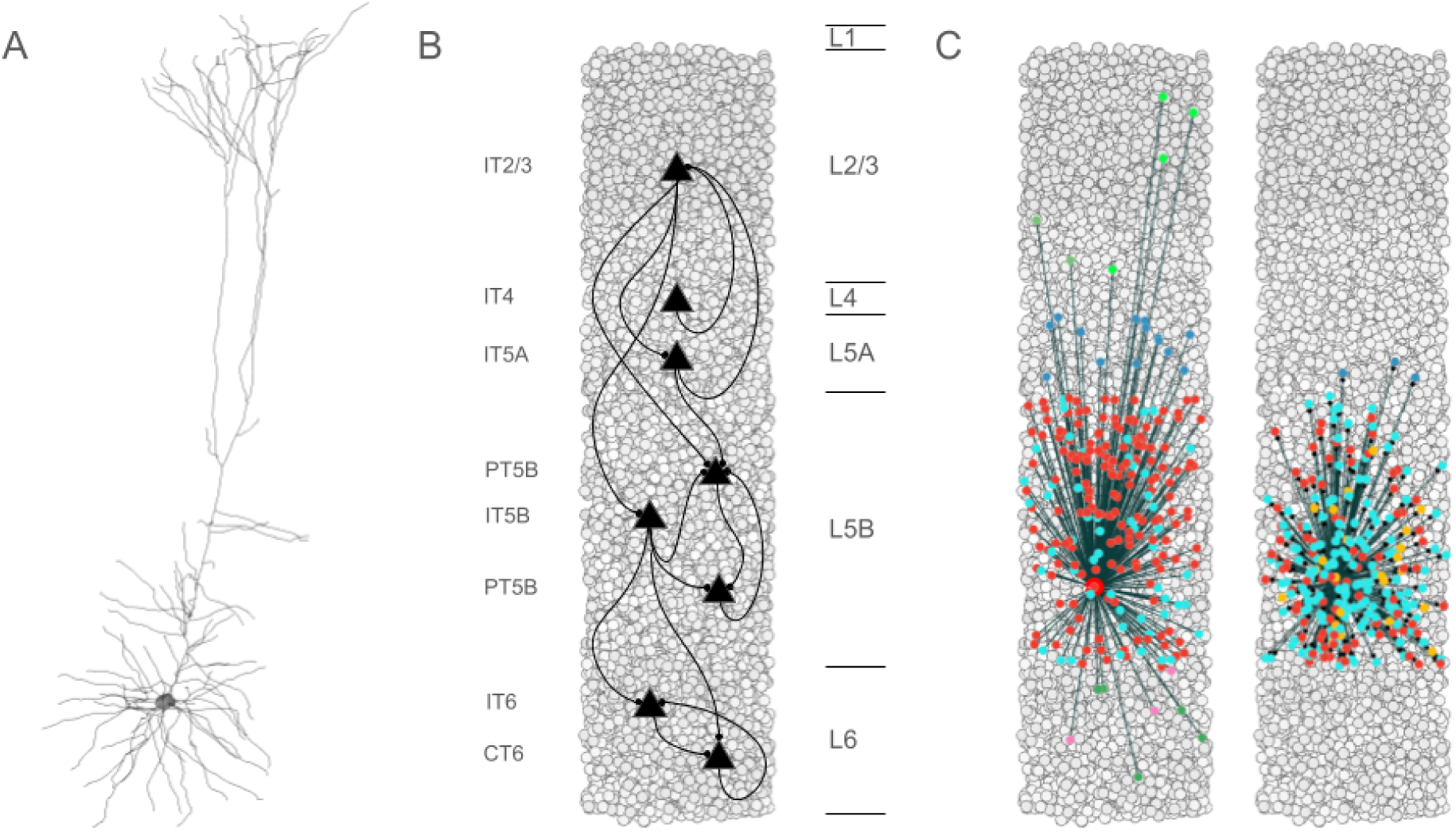
Simulated primary motor cortex (M1) is composed of 10,073 neurons. **A.** Layer 5 pyramidal tract neuron (PT5B) morphology had more than 700 compartments and multiple ion channel mechanisms. **B.** Pyramidal cell population connectivity across layers in a cylindrical column of M1 with 300 μm diameter and 1350 μm height (full cortical depth). **C.** Connectivity for a single PT5B neuron. Left: convergence from PT5B cells (red) and inhibitory neurons. Right: divergence to other PT5B cells (red) and inhibitory neurons at right. Inhibitory neuron color code -- L2/3 somatostatin-staining cell (SOM2/3) : light green; PV2/3 parvalbumin cell : dark green; PV5A : dark blue; SOM5B : orange; PV5B : light blue; SOM6: pink; PV6 : dark green.

### Control simulations

Comparison of rest and activated states in the control condition revealed dominant beta-band activity (∼20 Hz) in the rest state that transitioned to gamma-band activity (∼44 Hz) with the motor thalamic input changed to activated-state activity (Fig 2). In the rest state, each layer 5 pyramidal cell population (IT5A, IT5B, PT5B) showed different rates and durations of spiking activity with respect to each of the other populations during beta-band oscillations (Fig 2A). IT5A neurons were active from around the low point of PT5B activity, visible in the blue PT5B spike count histogram (SCH) at bottom of Fig 2A, and increased to around peak PT5B activity. IT5B neurons were primarily active during the increase in PT5B activity, from around the beginning of the PT5B activity duty cycle (see below), to around peak PT5B activity (Fig 2A, PT5B-SCH). In contrast, in the activated state, thalamic drive onto IT4 neurons lead to activation of PT5B and a decrease to zero activity in IT5A and IT5B (Fig 2B). The activated state showed greater gamma oscillation, seen in the LFP signal (Fig. 2AB, top; also see Fig 3B) as high frequency low-amplitude periodic deflections with intermittent periods of oscillatory activity at higher amplitude (∼44 Hz; Fig 2B). The beta frequency band in the LFP signal during rest and the gamma-band activity during the activated state were reflected in antiphase oscillations in their respective PT5B-SCH.

**Figure 2.**
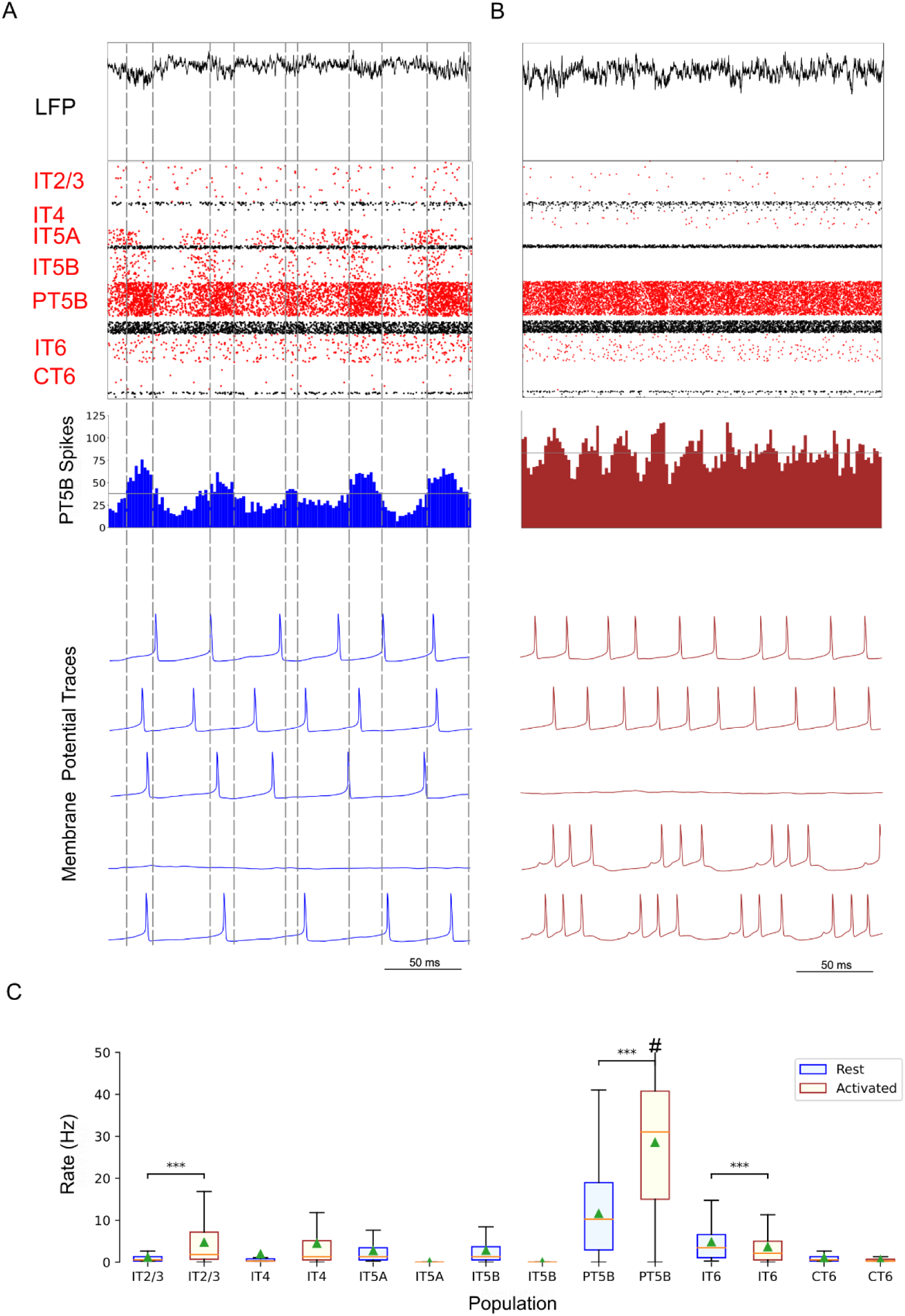
Control condition, simulated primary motor cortex (M1) activity, comparing resting state to movement state. **A.** Resting state (250 ms at midpoint of 4.3 s simulation) displayed dominant beta-band activity (∼20 Hz; particularly clear in the PT5B spike count histogram). Top-to-bottom: local field potential (LFP), raster plot (red excitatory; black inhibitory neurons), PT5B spike count histograms (2 ms bins; dashed lines at half height), and 5 randomly chosen PT5B voltage traces. **B.** Activated state showed change to high-frequency gamma-band activity (∼44 Hz). Same top-to-bottom as in A; bottom traces are the same neurons as in A. **C.** Statistically significant change was found for firing rates for excitatory cell types going from rest to activated: IT2/3 -- rest: 1.2±1.6 spikes/s; activated: 6.7±8.9 spikes/s; PT5B -- 11.7±9.4 vs 26.8±18.2 spikes/s; IT6 neurons -- 5.8±5.8 vs 3.5±5.2 spikes/s (mean, std dev, range shown; ***p<0.001; #: truncated maximum rate 73 spikes/s; calculated from the last 2.3 s of 4.3 s simulation to avoid initialization transients).

**Figure 3.**
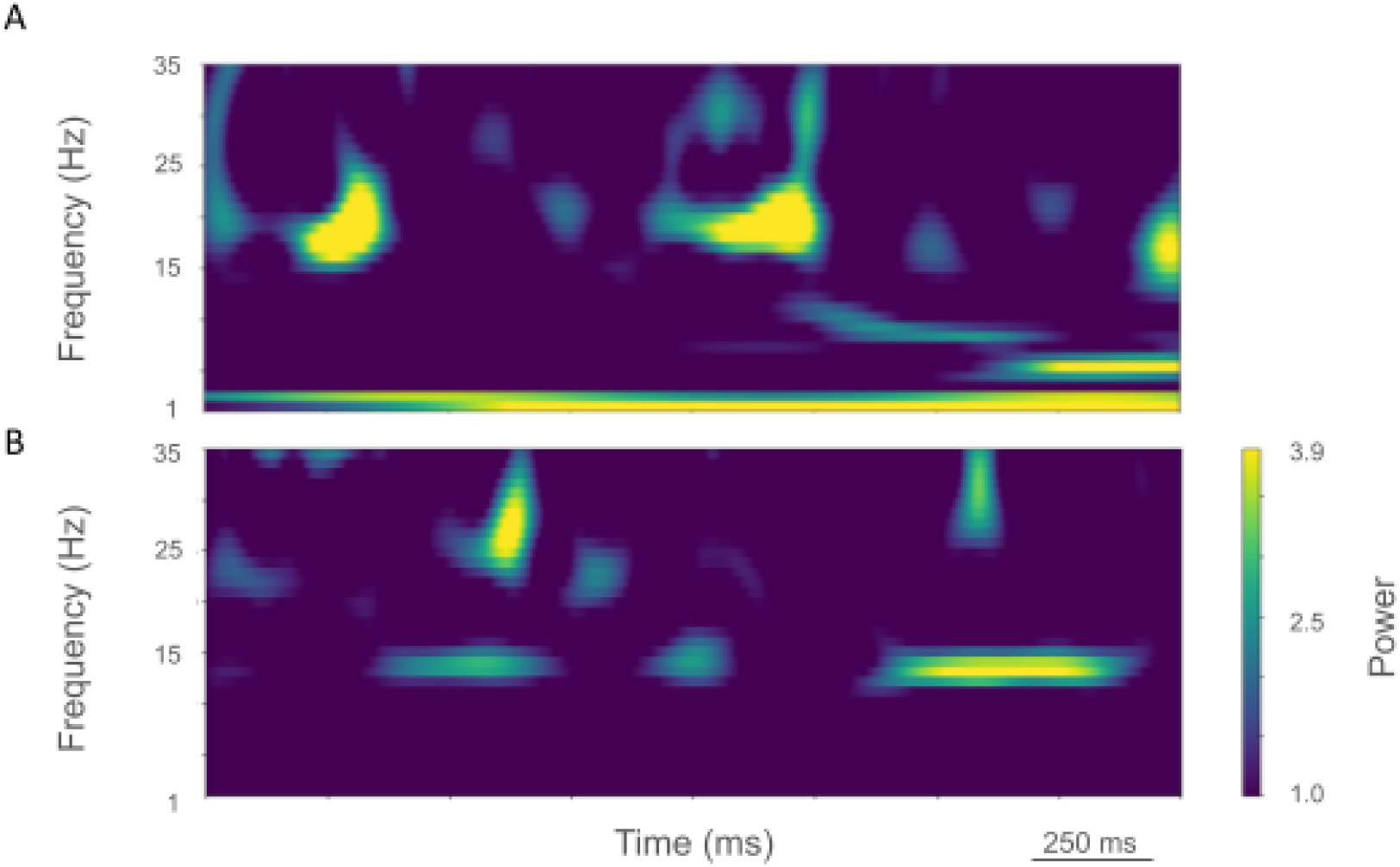
Time-resolved spectrograms of cortical LFPs in M1 in the control condition, under rest and activation states. **A.** The rest state simulation revealed continuous power around 2 Hz, as well as 15-25 Hz beta-band bursts occurring every 600 ms (burst rate 1.7 Hz). **B.** During the activated state the 2 Hz activity disappeared and beta bursts shifted to 15 Hz bursts ∼100-500 ms in duration or about 1.5 Hz. Beta-band power between 15-20 Hz was no longer visible. Higher frequency brief bursts of activity (∼50 ms duration) of 25-35 Hz occurred at about 1 Hz. (2.3 s of activity starting 2 s after simulation initiation. Color coding power in bar x 10^−5^).

PT5B neurons greatly increased their spiking overall, and changed their firing pattern in the activated state (Fig 2C). We analyzed the coefficient of variation (CV) of interspike intervals (ISI) observed across the PT5B neuron population to identify patterns of firing. The CV was found to be highly variable across the PT5B population during rest (0.59 +/− 0.37), indicating the presence of both regular (CV <= 0.5; 47.8%) and irregular activity (CV > 0.5; 52.2%). Using the same definitions, PT5B populations showed primarily irregular spiking in the activated state, again with high variability between neurons (overall CV 0.61 +/− 0.30; 25.7% CV <= 0.5; 74.3% CV > 0.5). Oscillations in subthreshold membrane potentials appeared frequently during the activated state (57.4% of PT5B neurons) but not during rest (Fig 2AB). Prominent excitatory postsynaptic potentials (EPSPs) appeared before the first spike in oscillatory traces in the activated state but not in the rest state or in non-oscillatory traces in the activated state. EPSPs appear in the activated state due to increased norepinephrine (NE) which reduces Ih conductance. A functional decrease in Ih results in increased input resistance, reduced sag, and eliminated resonance resulting in more robust temporal summation of EPSPs in PT5B neurons with reduced Ih.^60^

We explored oscillatory activity patterns further, identifying the oscillation period as beginning during increasing spike counts where the spike number was halfway between the minimum and maximum spike counts (Fig 2AB; horizontal gray line in PT5B-SCH gives half height; dashed vertical lines show half-height at rising and falling; distance along x-axis between 2 vertical lines is a single period). Each period was divided into high activity (above half-height) versus low activity (below) sections. We defined a duty cycle as the percentage of time during one oscillation period with relatively high activity (above half-height). The duty cycle was 31% during rest (Fig 2A) and 48% during activated state (Fig 2B; dashed vertical lines not shown).

Pyramidal tract neurons had the highest spike rate of any neuron population during the rest state, and increased further in the activated state (rest: 11.7±9.4 spikes/s; activated: 26.8±18.2 spikes/s; p<0.001;Fig 2C). In contrast, IT5A and IT5B firing rates decreased from 3 Hz (IT5A: 3.3 spikes/s; IT5B: 3.1 spikes/s) at rest, to 0 spikes/s in the activated state, and IT6 firing rates decreased from 5.8 spikes/s at rest to 3.5 spikes/s in the activated state (p<0.001). IT2/3 neurons significantly increased their firing rate (rest: 1.2 spikes/s; activated: 6.6 spikes/s; p<0.001). The IT4 neuron population firing rate increased from 2.6 Hz during rest to 4.7 Hz during the activated state.

The rest state simulation revealed virtually continuous power around 2 Hz, as well as 15-25 Hz beta-band bursts (Fig 3A), occurring every 600 ms (or with a burst-rate of 1.7 Hz). During the activated state the 2 Hz activity disappeared and beta bursts shifted to 15 Hz bursts ∼100-500 ms in duration or about 1.5 Hz. Beta-band power between 15-20 Hz was no longer visible (Fig 3B). Higher frequency brief bursts of activity (∼50 ms duration) 25-35 Hz occurred once every 1000 ms or about 1 Hz.

We looked for spikes that fired in the PT5B neuron population within a 1 ms time window of one another and considered those to be coincident spikes. The proportion of spikes that were coincident with that of other PT5B neurons within 1 ms was substantially higher in the activated M1, as compared to simulation of M1 at rest (Fig 4). PT5B spike coincident firing was observed in 1% of spikes at rest and increased to 3% during the activated state (Fig 4, cyan lines in A,C cyan + red in raster in B,D). PT5B coincident firing that exceeded those expected from frequency-matched random Poisson processes (significance at p<0.05; joint-surprise test^71^) was seen during rest in 100-275 ms duration clusters (red squares in Fig 4B). The mean periodic activity during significant coincident events (red bands) was 19 Hz as measured during each period (red band) of significant synchrony. The range of periodic activity across individual bands of significant synchrony was 15-24 Hz. During the activated state, more frequent and denser clusters or periods of synchrony were seen than during the rest state, with durations of 125-250 ms (red squares in Fig 4D). The total duration of significant synchronous activity increased from 0.7 s out of 4.0 s (17.5%) at rest to 1.1 s out of 4.0 s (27.5%).

**Figure 4.**
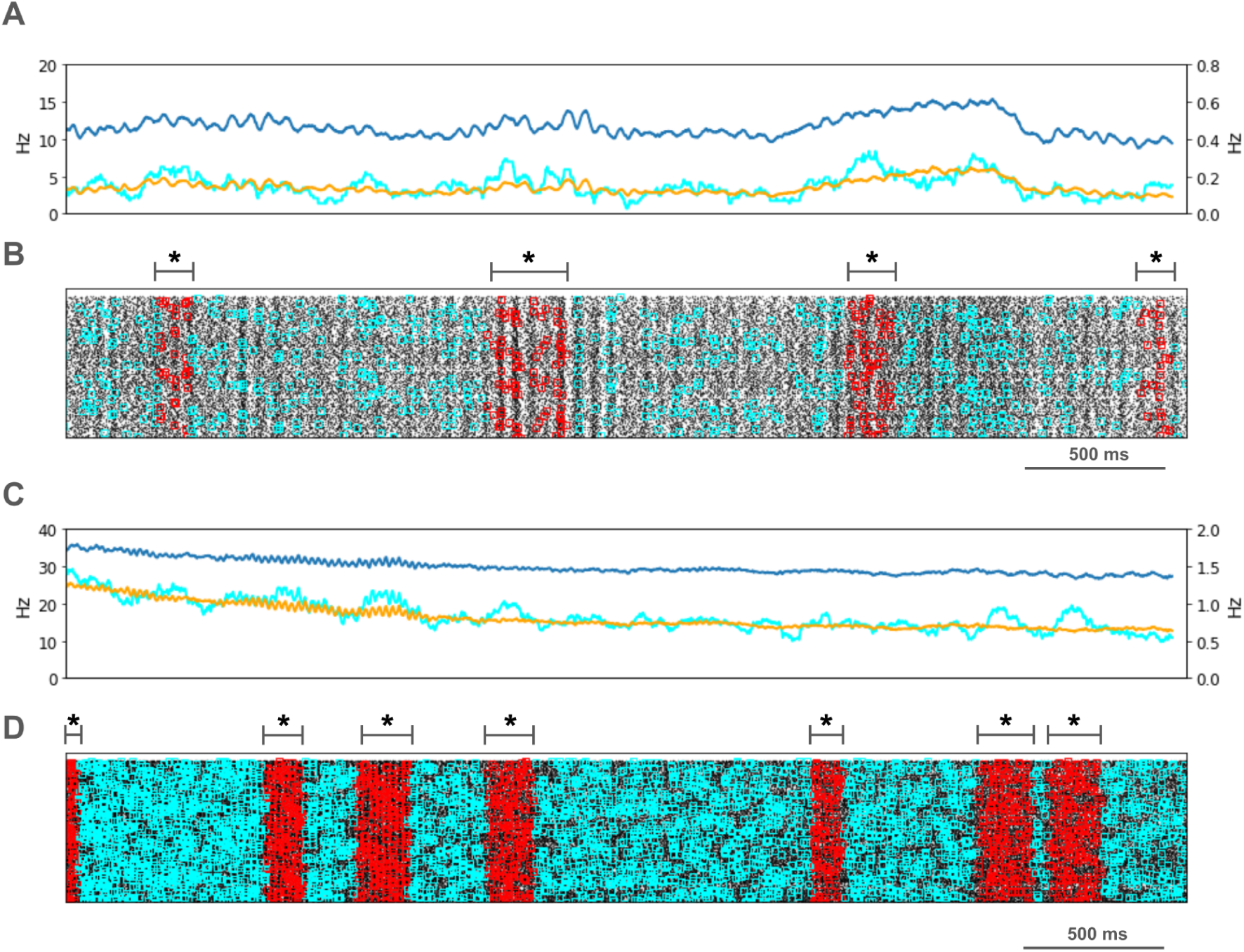
PT5B coincident firing during control condition. **A.** Resting state: coincidence rates (cyan; left y-axis) exceed expected (orange) which tracks overall rate (blue; right y-axis). **B.** Resting state: coincident events (cyan) and periods of significant coincident events (* p < 0.05; joint-surprise) with individual events in red.^71^ Significant coincident events appeared in 100-275 ms duration clusters (red bands). The total duration of significant synchronous activity during rest was 0.7 s out of 4.0 s (17.5%). **C.** Activated state: coincidence rates (cyan; left y-axis) exceed expected (orange) which tracks overall rate (blue; right y-axis). **D.** Activated state: coincident events (cyan) and periods of significant coincident events (* p < 0.05; joint-surprise^71^; 100ms firing rate window^72^) with individual events in red. Significant coincident events appeared in 125-250 ms duration clusters (red bands). The total duration of significant synchronous activity during the activated state was 1.1 s of 4.0 s (27.5%).

### Parkinsonian condition

Comparison of the simulated rest and activated states in the parkinsonian condition revealed the presence of focused 15 Hz beta-band activity in the rest state that transitioned to gamma-band activity (∼43 Hz) in the activated state (Fig 5). In the rest state, the layer 5 pyramidal cell populations (IT5A, IT5B, PT5B) showed beta band oscillations with slightly different phases (Fig 5A, raster diagrams): IT5A and IT5B neurons were active during the leading phases in the PT5B-SCH oscillation (Fig 5A). In contrast, in the activated state, the thalamic drive onto IT4 neurons led to activation of PT5B neurons, and stopped activity in IT5A and IT5B. The activated state showed greater gamma oscillation in the simulated LFP signal (Fig 5B), as higher frequency low-amplitude periodic deflections with intermittent higher amplitude excursions (∼44 Hz). The beta frequency band in the LFP signal during rest and the gamma-band activity during the activated state were reflected in antiphase oscillations in their respective PT5B-SCH.

**Figure 5.**
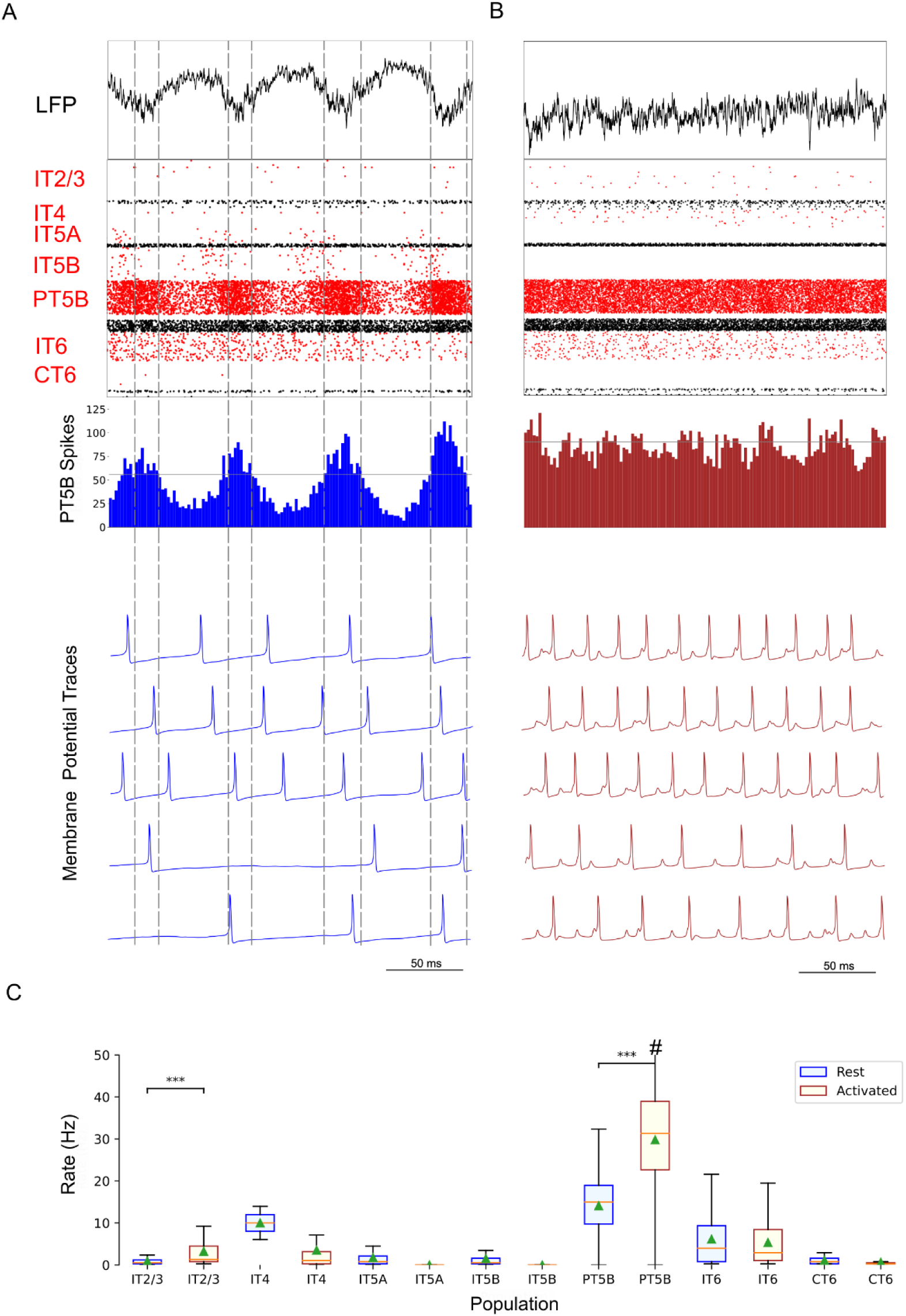
Parkinsonian condition, simulated primary motor cortex (M1) activity, comparing resting state to movement state. **A.** Resting state (250 ms shown at midpoint of 4.3 s simulation) displayed focused 15 Hz beta-band activity. Top-to-bottom: local field potential (LFP), raster plot (red excitatory; black inhibitory neurons), PT5B spike count histograms (2 ms bins; dashed lines at half height), and 5 randomly chosen PT5B voltage traces. **B.** Activated state showed change to high-frequency gamma-band activity (∼43 Hz). Same top-to-bottom as in A; bottom traces are the same neurons as in A. **C.** Statistically significant change was found for firing rates for excitatory cell types going from rest to activated: IT2/3 -- rest: 0.78±1.1 spikes/s; activated: 4.3±7.1 spikes/s; PT5B -- rest: 14.2±7.1 vs 29.7±12.7 spikes/s (mean, std dev, range; ***p<0.001; #: truncated maximum rate 65.7 spikes/s; calculated from the last 2.3 s of 4.3 s simulation to avoid initialization transients).

PT5B neurons both greatly increased spiking and changed firing pattern in the activated state (Fig 5C). The CV of the PT5B neuron population ISIs analysis showed a peak signifying regular activity (CV of 0.34 +/− 0.27) but with a large enough standard deviation to include irregular activity in the PT5B population during rest: regular (80.6% CV <= 0.5) and irregular activity (19.4% CV > 0.5). In the activated state, PT5B population analysis showed regular spiking that included a small number of irregular spiking neurons (CV 0.18 +/− 0.19; 93.8% CV <= 0.5 regular; 6.2% CV > 0.5 irregular). Subthreshold EPSPs were clearly evident during the activated state but not during resting (Fig 5AB). Activated state EPSPs appear larger in the parkinsonian in contrast with the control condition due to the greater driving force on EPSP since, on average, the membrane potential is more negative in the parkinsonian condition with its reduced excitability. No subthreshold membrane potential oscillations were observed in the parkinsonian rest or active states. The duty cycle was 29% during the parkinsonian rest state (Fig 5A) and 25% during activated state (Fig 5B; dashed vertical lines not shown).

Pyramidal tract neurons had the highest spike rate of any neuronal population during the rest and activated states in the parkinsonian condition (rest: 14.2±7.1 spikes/s; activated: 29.7±12.7 spikes/s; p<0.001; Fig 5C). IT5A and IT5B firing rates decreased from 2.2 and 1.2 spikes/s, respectively, at rest to 0 spikes/s in the activated state. The firing rates of IT6 neurons did not change (rest and activated: 6.0 spikes/s). In contrast, superficial layer IT2/3 neurons significantly increased their average firing rates with activation (rest: 0.8 spikes/s; activated: 4.3 spikes/s; p<0.001).

LFP spectrograms of resting oscillatory power displayed continuous high-power 15 Hz beta-band power with bursts of power in the ∼25-35 Hz (∼50-250 ms duration) occurring about once every 200 ms (∼5.0 Hz; Fig 6A). In the activated state, ∼20-35 Hz bursts ∼25-100 ms in duration were seen that occurred every ∼50-100 ms (∼10-20 Hz; Fig 6B).

**Figure 6.**
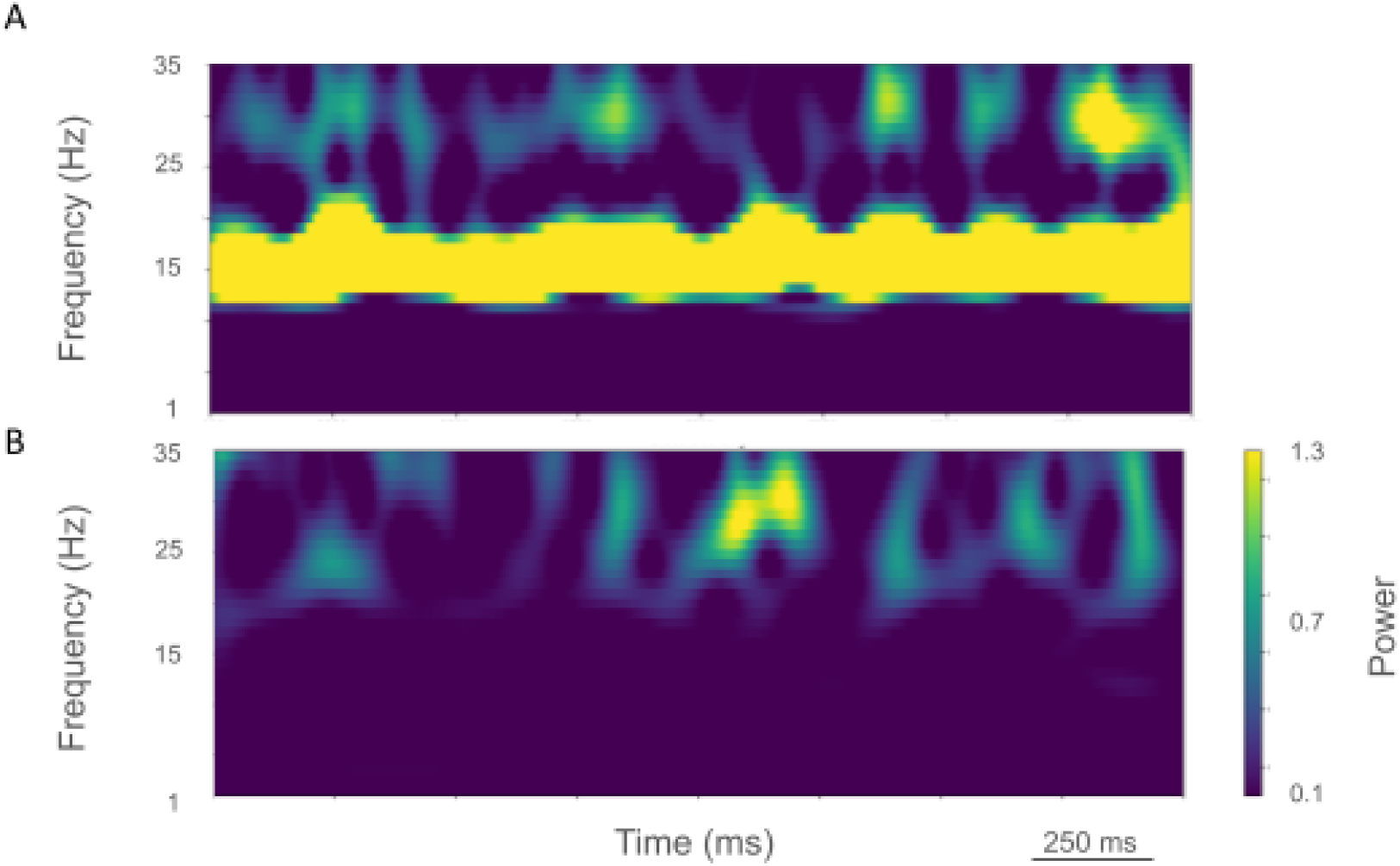
Time-resolved spectrograms of cortical LFPs in M1 in the parkinsonian condition, under rest and activation states. **A.** The rest state simulation revealed a continuous high-power 15 Hz beta-band with bursts of power in the ∼25-35 Hz (∼50-250 ms duration) occurring about once every 200 ms (∼5.0 Hz). **B.** During the activated state, ∼20-35 Hz bursts of ∼25-100 ms duration occurred every ∼50-100 ms (∼10-20 Hz) (2.3 s of activity starting 2 s after simulation initiation. Color coding power in bar x 10^−4^).

PT5B spike coincident firing (2 or more spikes firing in 1ms) was observed in 2% of spikes at rest and increased to 3% during the activated state (Fig 7, cyan lines in A, C cyan + red in raster in B, D). PT5B coincident firing that exceeded those expected from frequency-matched random Poisson processes (p<0.05; joint-surprise test^71^) was seen during rest in 400-1,100 ms duration clusters with a periodic structure within each, consisting of 6 to 16 periods (red squares in Fig 7B). The mean periodic activity during significant coincident events (red bands) was 15 Hz as measured during each period (red band) of significant synchrony. The range of periodic activity across individual bands of significant synchrony was 14-15 Hz. During the activated state, the significant PT5B coincident firing (p<0.05) occurred in epochs of dense synchronous activity, 100-350 ms in duration (red squares in Fig 7D). The total duration of significant synchronous activity during rest was 3.1 s out of 4.0 s (77.5%). During the activated state, 1.4 s of 4.0 s (35.0%), was significantly synchronous.

**Figure 7.**
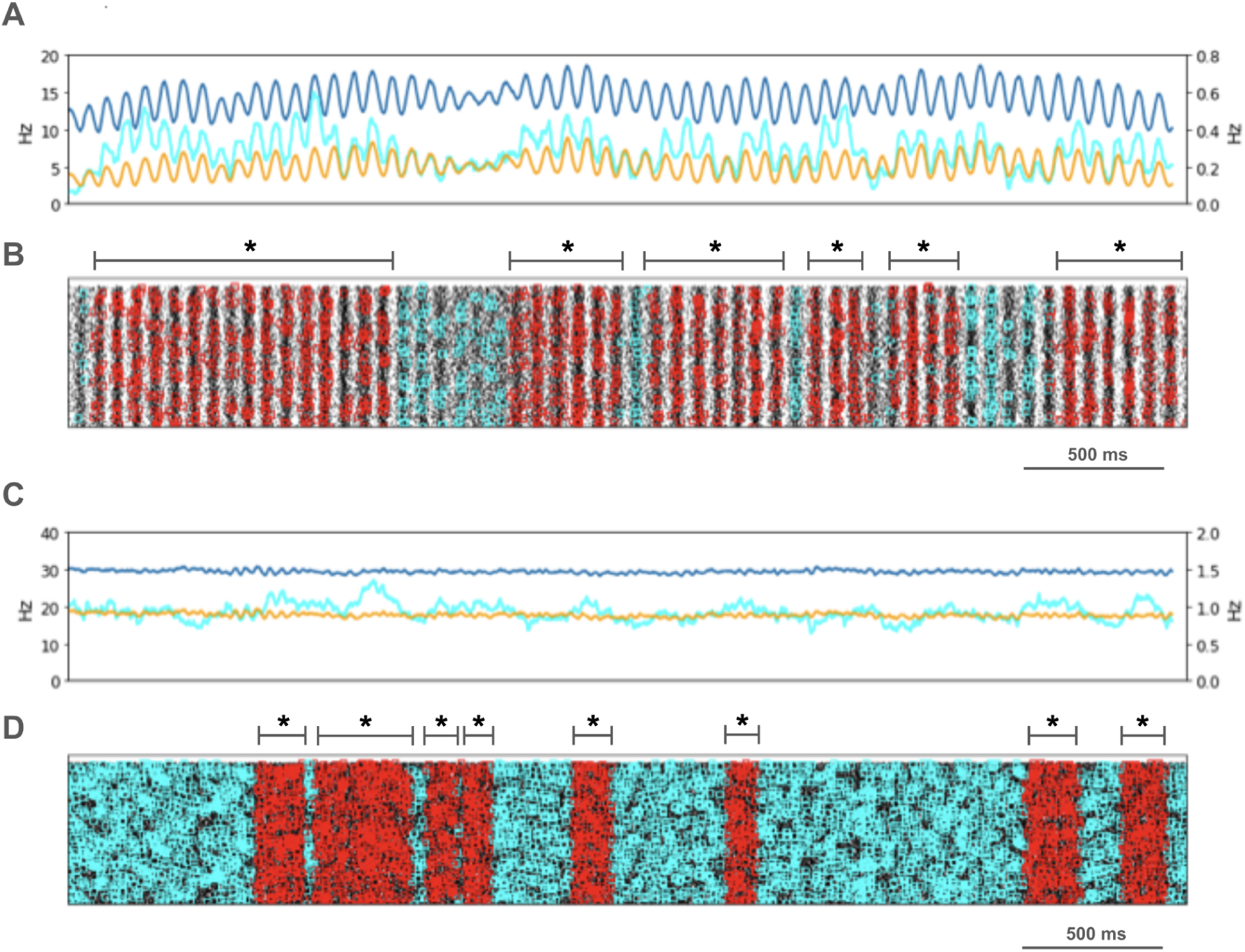
PT5B coincident firing during parkinsonian condition. **A.** Resting state: coincidence rates (cyan; left y-axis) exceed expected (orange) which tracks overall rate (blue; right y-axis). **B.** Resting state: coincident events (cyan) and periods of significant coincident events (* p < 0.05; joint-surprise) with individual events in red. Significant coincident events appeared in 400-1,100 ms duration clusters (red bands). The total duration of significant synchronous activity during rest was 3.1 s out of 4.0 s (77.5%). **C.** Activated state: coincidence rates (cyan; left y-axis) exceed expected (orange) which tracks overall rate (blue; right y-axis). **D**. Activated: coincident events (cyan) and periods of significant coincident events (* p < 0.05; joint-surprise^71^; 100ms firing rate window^72^) with individual events in red. Significant coincident events appeared in 100-350 ms duration clusters (red bands). The total duration of significant synchronous activity during the activated state was 1.4 s of 4.0 s (35.0%).

### Control vs parkinsonian comparison

The most substantial changes in firing between conditions were seen in the simulated rest state, where there was a significant increase in PT5B firing rate and a decrease in firing rate variability in the parkinsonian condition as compared with control condition (control: 11.7±9.4 spikes/s; parkinsonian: 14.2±7.1 spikes/s; p<0.001). There was also an increase in IT4 firing rate (control: 2.6 spikes/s; parkinsonian: 14.1 spikes/s; p<0.05) and a decrease in IT5A firing rate (control: 3.3 spikes/s; parkinsonian: 2.2 spikes/s; p<0.01).

Activated (movement-associated) cortical dynamics also showed a significant increase in PT5B firing rate and a decrease in firing rate variability in the parkinsonian condition as compared with control condition (control: 26.8±18.2 spikes/s; parkinsonian: 29.7±12.7 spikes/s; p<0.001). The firing rate of IT6 neurons also increased in the parkinsonian condition (control: 3.5 spikes/s; parkinsonian: 6.0 spikes/s; p<0.001).

## Discussion

We performed a series of simulations based on an isolated cortical change, reduced PT5B excitability, found in parksionian mouse models.^19,73^ In the resting state, reduced PT5B neuron excitability resulted in a paradoxical increase in PT5B firing in the network condition, as well as an increase in beta oscillatory power with reduced frequency, increased PT5B spike synchrony, and firing rate shifts in other cell populations. Parkinsonism-associated changes were less marked in the activated (movement) state, where we found no significant change in PT5B population firing rate but did see a change in PT5B activity pattern with an increase in power of 20-35 Hz activity, comparable to what is seen in patients.^74,75^

Shifts in dominant frequency activity were prominent in resting versus activated data. At rest we observed 2 Hz oscillations and 15-20 Hz beta-band bursts in LFP signals once every ∼0.6 s. In contrast, during the activated state, only the lowest frequency beta bursts of ∼15 Hz remained, with high beta and low gamma bursts appearing in the 25-35 Hz band. These findings are in line with previous results indicating that beta-rhythms are known to desynchronize during voluntary movement.^76,77^ Within this context, it is interesting that our change to produce the activated state consisted of only two changes: 1) an increase in random spikes from thalamus from 5 Hz to 10 Hz, and 2) a decrease in H current in PT5B neurons. These relatively small changes were sufficient to result in decreased power across most beta band rhythm frequencies in M1.

The parkinsonian condition was marked by the appearance of vigorous 25-35 Hz band bursts in both the resting and activation states. In the resting state, an order of magnitude increase in beta power was observed around 15 Hz. With activation, the high-beta and low-gamma oscillation bursts (25-35 Hz) extended to lower frequencies around 20 Hz. These changes were produced by a decrease of the excitability of cortical neurons, which paradoxically resulted in the increased PT5B spiking rates within the context of the simulated M1 circuit.

In healthy individuals, an early hypothesis was that beta-oscillatory bursts in M1 seen by EEG or electrocorticogram were a marker of an “idling state” before the desynchronization with movement.^76–78^ A more recent “status quo” hypothesis states that beta band activity has a stabilizing effect, signifying active processes that promote existing motor set while suppressing the neuronal processing of new movements.^77,79^ The frequency of beta oscillations is strongly coupled with the dopamine tone in monkeys and humans.^78,80^ Lesions of midbrain dopaminergic neurons in animals lead to an increase in beta-frequency oscillatory activity in the basal ganglia, M1,^16^ subthalamic nucleus, and globus pallidus.

Many studies have reported an increase in beta band power or in beta-oscillatory bursts in the basal ganglia, thalamus, or cortex of patients with PD or in animal models of dopamine loss.^18,80^ A connection between DA loss and beta activity was demonstrated in rodents when deep brain stimulation in parkinsonian rats destroyed the M1 dominance of beta rhythms and restored motor control.^81^ In addition, a recent study demonstrated beta oscillations in the basal ganglia of parkin knockout mice.^82^

The overall amount of beta-band power in LFP signals reflects the average of beta burst^83^ activity during the period that was examined. Although not universally agreed upon,^84^ there may be several key differences between the patterns of beta burst activity in healthy and parkinsonian subjects. Thus, the timing and duration of beta power bursts is highly variable in healthy subjects, while the variability is much lower in patients with severe PD. Further, in contrast to findings in healthy subjects, beta bursts in M1 in parkinsonian individuals are often unusually long (even greater than 100 ms duration^83^). It has been suggested that the brevity of beta bursts in the healthy state could be critical to normal beta-band function.^83^

Waveform features of beta oscillations in LFP (or electrocorticogram) signals may reflect synchronous excitatory synaptic inputs onto cortical pyramidal neurons. Beta-band oscillations in M1 in parkinsonian patients have sharp, asymmetric, nonsinusoidal features that are correlated with beta-high gamma phase-amplitude coupling. The observation of sharp beta oscillations in PD M1 due to synchrony of synaptic activity has been hypothesized to be due to increases in beta synchrony in the basal ganglia. We observed changes in synchrony and significant increases in beta oscillations in simulated M1 with only small changes in potassium and sodium currents and only in PT5B neurons. Observed changes were intracortical changes in activity patterns due to M1 circuitry and biophysics.

Oscillations in the brain provide an effective means to control the timing of neuronal firing. Cortical neurons support highly precise and reliable spike times to naturalistic fluctuating inputs^85^ and are good detectors of correlated activity.^86^ Oscillations can temporally coordinate information transfer and support spike-timing dependent plasticity.

Significant increases in synchronous neural activity in M1 is consistently observed in PD^87^ or in animal models of the parkinsonian condition, including an increase in the concurrence of beta bursts.^84^ We found a substantial increase in synchronous spiking in our simulations of the parkinsonian rest state.

Major limitations of this study are the limitations that are inherent in all modeling studies—we necessarily made choices as to what to include and what to leave out. Many parameters are not considered since they have not been studied experimentally or cannot currently be studied in detail (this includes, e.g., functions of dendritic spines). In particular, (1) we did not consider interneuron populations other than PV and SOM cells; (2) we did not consider the cortical effects of dopamine; (3) we modeled inhibitory neurons with three compartments; (4) we have incomplete models for the distribution of voltage- and calcium-sensitive dendritic channels in pyramidal dendrites. We qualitatively matched PT5B data in control and 6-OHDA conditions by manually modifying conductances. Finally, our simulations showed increased PT5B spiking rates within the context of the simulated M1 circuit. However, reports from *in vivo* MPTP-macaque recordings showed marked decreased PT5B firing rates in parkinsonian animals.^16–18^ This discrepancy can be resolved by going outside of the M1 circuit. A recent study showed reduced thalamocortical synaptic strength onto the layer 5 dendrites of PT5B neurons in the 6-OHDA mouse M1.^73^ This reduced excitatory current will alter the overall activity balance in PT5B and will be evaluated in future simulations.

A small local change in cortical PT5B neuron excitability sufficed to induce changes in cortical oscillations that resemble the parkinsonian condition. This change in the experimental animal occurred as a consequence of a treatment (6-OHDA) with effects on basal ganglia activity as well as on cortex directly. The combined consequences in this parkinsonian model, as well as in PD itself, will involve combined effects at several sites. The current study demonstrates that the effects of the subcortical dopamine loss is not solely due to the transmission of abnormal subcortical signals to the cerebral cortex, but will be due to local cortical effects as well. Decrease in PT5B excitability is sufficient change to produce abnormal M1 oscillatory activities which will, in turn, alter basal ganglia activity, given that all of these are components of the larger coupled oscillator.

## Methods

We utilized our previously developed network simulations of the mouse M1,^21^ written using the NEURON/NetPyNE simulation platform.^22–24^ NEURON version 8.2.2 (https://www.neuron.yale.edu, RRID:SCR_005393) and NetPyNE version 1.0.5 (https://netpyne.org, RRID:SCR_014758) are open source software. The fully-commented simulations of this paper are available at GitHub (https://github.com/suny-downstate-medical-center/pd_m1_6ohda) and through Zenodo (http://doi.org/10.5281/zenodo.12399983) under CC BY 4.0 License, freely available for modification, redistribution and research use by the community (feel free to email authors with questions).^25–27^ All resources are listed in the Supplementary Data Key Resource Table (Supplementary Table 1).

To demonstrate robustness, we ran 4 sets of 4 simulations (Supplementary Table 2), each set including the control and parkinsonian conditions in rest state and in the activated (movement) state. Each set used unique random seeds for connectivity, stimulation, and neuron location. In this paper we used the experiment set sM1_12-12-2023 as a typical example. Analyses of the 3 other sets of experiments are shown in Supplementary Table 2 and a typical example set of Supplementary Figures 2-5.

We compared the control “healthy” condition with a model of the parkinsonian condition based on our co-authors physiology of the 6-hydroxydopamine treated, dopamine-depleted mouse (6-OHDA mouse).^19^ These *in vitro* data showed changes in excitability in PT5B but not in IT neurons in 6-OHDA mouse. PT5B spike count decreased 64% on average across current injections (Fig. 1D of Chen et al 2021^19^; following figure references also from Chen 2021); BK (Figs. 6, 7) and NaT (Fig. 5) currents changed (Fig. 4). Therefore, we first increased BK conductance which, as expected, hyperpolarized PT5B membrane potential, reducing PT5B excitability since further from threshold. However, the slope of the current-frequency (I-f) curve was too shallow so we increased NaT to more closely match (Supplementary Figure 1 of present paper; Supplementary Table 3).

### Morphology and physiology of neuron classes

Seven excitatory pyramidal cell types and two interneuron cell types were simulated in the network (Fig 1). Our detailed multicompartment model for PT5B was based on prior layer 5B PT5B *in vitro* electrophysiological studies of the responses of these cells to somatic current injections.^28^ We optimized the parameters of the Hodgkin-Huxley neuron model ionic channels within a range of values constrained by the literature. The models included subtypes of excitatory pyramidal cells: PT5B, intratelencephalic (IT) and corticothalamic (CT) neurons (Fig 1B), inhibitory model neurons were: parvalbumin- (PV-) and somatostatin- (SOM-) containing interneurons. The Layer 5 (L5) PT5B and IT cell model morphologies had 706 and 325 compartments, respectively, digitally reconstructed from 3D microscopy images. Morphologies are available via NeuroMorpho.Org (RRID:SCR_002145; archive name “Suter_Shepherd”). We employed existing simplified 3-compartment (soma, axon, dendrite) models and increased their dendritic length to better match the average f-I slope and rheobase experimental values of cortical basket (PV) and Martinotti (SOM) cells (Neuroelectro online database). We implemented models for two major classes of GABAergic interneurons: parvalbumin-expressing fast-spiking (PV) and somatostatin-expressing low-threshold spiking neurons (SOM). In the text and figures, the abbreviated names are followed by the corresponding layer number including 2/3 (layers 2,3 together) and 5A vs 5B. Although M1 is classified as agranular cortex, we included layer 4 (L4) cells, based on previous experimental studies.^29^

In the M1 simulation, neuronal activities were driven by ascending input from ventromedial thalamus (VM) to layer 2/3 (L2/3), L4, L5A neurons and also from the ventrolateral thalamus (VL) onto L4, L5B; from primary and secondary somatosensory cortices (S1 and S2) to L2/3, L5A; from contralateral primary (cM1) and ipsilateral secondary motor cortices (M2) to L5B, L6; and from orbital cortex (OC) to L6 as described in the original model.^21^ Each input region consisted of a population of 1000 spike-generators (NEURON VecStims) that generated independent random Poisson spike trains (based on experimental background activity: VL 0-2.5 Hz; VM of 0-5 Hz; S1, S2, OC 0-5 Hz; cM1, M2 0-2.5 Hz).

The M1 model in the current study was identical to M1 validated in the Dura-Bernal et al. (2023) paper except in the current study control PT5B currents were slightly modified (NaT, and BK; Supplementary Table 2) so that their response properties were more similar to the mean control PT5B neurons reported by Chen et al. (2021). In the simulated parkinsonian mouse M1 PT5B currents (BK and NaT) were modified so that they showed a 64% decrease in excitability as reported by Chen et al. (2021). In vitro data pointed to changes in BK and NaT currents in the PT5B neurons of parkinsonian mice. Increasing BK current moved the PT5B neuron’s membrane potential away from threshold (more negative) but slope of the current-frequency curve was too shallow. A small increase in NaT current increased the slope of the current-frequency curve to more closely match the values seen in vitro (Supplementary Figure 1).

### Microcircuit composition: neuron locations, densities and ratios

We modeled a cylindrical volume of the mouse M1 cortical microcircuit with a 300 μm diameter and 1350 μm height (cortical depth) at full neuronal density for a total of 10,073 neurons. Cylinder diameter was chosen to approximately match the horizontal dendritic span of a corticospinal neuron located at the center, consistent with the approach used in the Human Brain Project model of the rat S1 microcircuit.^30^ Mouse cortical depth and boundaries for layers 2/3, 4, 5A, 5B and 6 were based on published experimental data.^29,31,32^ Although traditionally M1 has been considered an agranular area lacking L4, M1 pyramidal neurons were recently identified with the expected prototypical physiological, morphological, and wiring properties of L4 neurons ^29,33,34^ and therefore were incorporated in the model.

Cell classes present in each layer were determined based on mouse M1 studies.^29,32,35–39^ IT cell populations were present in all layers, whereas the PT5B cell population was confined to layer 5B, and the CT cell population only occupied layer 6. SOM and PV interneuron populations were distributed in each layer. Neuronal densities (neurons per mm^3^) for each layer were taken from a histological and imaging study of mouse agranular cortex.^40^ The proportion of excitatory to inhibitory neurons per layer was obtained from mouse S1 data.^41^ The proportion of IT to PT5B and IT to CT cells in layers 5B and 6, respectively, were both estimated as 1:1.^35,36,42^ The ratio of PV to SOM neurons per layer was estimated as 2:1 based on mouse M1 and S1 studies.^43,44^ Since data for M1 layer 4 was not available, interneuron populations labeled PV5A and SOM5A occupy both layers 4 and 5A. The number of cells for each population was calculated based on the modeled cylinder dimensions, layer boundaries and neuronal proportions and densities per layer.

### Local connectivity

Wiring was identical to that in our previous published studies of M1.^21,45^ In brief, data from multiple studies were used to provide type-to-type location connectivity based on mapping studies at 100 μm spatial resolution, well beyond traditional layer-based connectivity resolution: 1. E>E (E, excitatory neurons: IT, PT5B,CT) based on collaborator papers.^29,31,32,42^ Following previous publications ^41,46^ we defined S_con_ = p_con_ × v_con_ (subscript *con* is connection; strength (S_con_) between 2 types is product of probability p_con_ × unitary somatic EPSP amplitude v_con_, normalized based on NEURON connection weight of an excitatory synaptic input to generate a somatic EPSP of 0.5 mV at each neuron segment. For morphologically detailed cells (layer 5A IT and layer 5B PT5B), the number of synaptic contacts per unitary connection (or simply, synapses per connection) was set to five, estimated from mouse M1^47^ and rat S1.^30,48^ For the remaining cell models, all with six compartments or less, a single synapse per connection was used.

We adapted E>I (PV, SOM) strengths from studies by our collaborators,^42,49^ using their layer-based descriptions. Probability of connection decayed exponentially with the distance between the pre- and post-synaptic cell bodies with length constant of 100 μm.^50,51^ We introduced a correction factor to the distance-dependent connectivity measures to avoid border effect, i.e., cells near the modeled volume edges receiving less or weaker connections than those in the center. We rederived a full population to population connectivity on a layer basis to integrate the more location specific E>E with lower-resolution E>I connectivity.

Excitatory synapses colocalized AMPA (rise, decay τ: 0.05, 5.3 ms), NMDA (15, 150 ms) receptors: 1:1 ratio, both with E_rev_ = 0 mV 1.0,^52^ NMDA conductance scaled by 1/(1 + 0.28 · exp (−0.062 · V)); for Mg effect with [Mg] = 1mM.^53^ Inhibitory synapses from SOM to excitatory neurons consisted of a slow GABAA receptor (rise, decay τ: 2, 100 ms) and GABAB receptor, in a 90% to 10% proportion; synapses from SOM to inhibitory neurons only included the slow GABAA receptor; and synapses from PV to other neurons consisted of a fast GABAA receptor (rise, decay τ: 0.07, 18.2). The reversal potential was −80 mV for GABAA and −95 mV for GABAB. The GABAB synapse was modeled using second messenger connectivity to a G protein-coupled inwardly-rectifying potassium channel (GIRK).^54^ The remaining synapses were modeled with a double-exponential mechanism.

Connection delays were estimated as 2 ms plus a variable delay depending on the distance between the pre- and postsynaptic cell bodies assuming a propagation speed of 0.5 m/s.

### Dendritic distribution of synaptic inputs

Experimental evidence demonstrates the location of synapses along dendritic trees follows very specific patterns of organization that depend on the brain region, cell type and cortical depth ^55,56^; these are likely to result in important functional effects.^57–59^ Synaptic locations were automatically calculated for each cell based on its morphology and the pre- and postsynaptic cell type-specific radial synaptic density function. Synaptic inputs from PV to excitatory cells were located perisomatically (50 μm around soma); SOM inputs targeted apical dendrites of excitatory neurons ^38,44^; and all inputs to PV and SOM cells targeted apical dendrites. For projections where no data synaptic distribution data was available – IT/CT to IT/CT cells – we assumed a uniform dendritic length distribution.

### Resting and activated states

The resting (quiet wakefulness) and activated (movement) states were simulated as validated by Dura-Burnal et al. (2023). In particular, the activated state was simulated by increasing thalamic input activity from motor thalamus to 0-10 Hz (uniform distribution), and reducing hyperpolarization-activated current (Ih) conductance to 25% in PT5B neurons, to simulate a high level of norepinephrine input from the locus coeruleus (LC).^21,60^ The other inputs continued to provide unchanged drive.

### Identifying synchronous spiking

We used unitary event analysis for identifying synchronous spiking significantly above the expected number of synchronous spikes for the neuron population size and firing rates.^61^

### Local field potential (LFP)

We used the local field potential (LFP) method built into the NetPyNE framework. LFP was calculated at each simulated electrode using the line source approximation,^62,63^ which is based on the sum of the membrane current source generated at each cell segment divided by the distance between the segment and the electrode. The calculation assumes that the electric conductivity (sigma = 0.3 mS/mm) and permittivity of the extracellular medium are constant everywhere and do not depend on frequency. We collected LFP signals from extracellular electrodes located at multiple depths within the M1 simulation.

### Duty cycle

Duty cycle is a common concept in electrical engineering and electronics and is defined as the ratio of time a load or circuit is on compared to the time the load or circuit is off. The use of the term has a rich history in neurophysiology ^64–66^ and in motor system neurophysiology in particular ^67–70^ where duty cycle often refers to the proportion of time the neuron or neural circuit is actively firing within a given period but can refer to other on-off cycles such as contracted versus relaxed muscle tissue. Duty cycle has also been defined as on when a threshold is achieved and otherwise off if not.^66^

In the current study, duty cycle is defined as on when 50% activity is reached from the trough (minimum activity) to the peak (maximum activity) during one period as measured in a spike histogram. A period is defined as the duration from the time increasing activity crosses the 50% activity threshold to the time that decreasing activity crosses the same threshold.

## Data Availability

The main and supplemental data generated in this study are freely available in NWB format in DANDI Archives (RRID:SCR_017571) https://dandiarchive.org/dandiset/001444 under the CC BY 4.0 License.

## Code Availability

The code used to generate data and carry out this study is openly available in GitHub https://github.com/suny-downstate-medical-center/pd_m1_6ohda and through Zenodo http://doi.org/10.5281/zenodo.12399983 under the CC BY 4.0 License.

## Acknowledgments

This research was funded by Aligning Science Across Parkinson’s [ASAP-020572] through the Michael J. Fox Foundation for Parkinson’s Research (MJFF) and by the National Institute of Neurological Disorders and Stroke (grant#: R01NS121371). This work used Expanse at San Diego Supercomputing Center through allocation IBN140002 from the Advanced Cyberinfrastructure Coordination Ecosystem: Services & Support (ACCESS) program, which is supported by National Science Foundation grants #2138259, #2138286, #2138307, #2137603, and #2138296. For the purpose of open access, the author has applied a CC BY public copyright license to all Author Accepted Manuscripts arising from this submission.

## Author Contributions

D.W.D. and W.W.L. contributed to the conception and design of the study; D.W.D., L.C., H.y.C., and W.W.L. contributed to the acquisition and analysis of data; D.W.D. and W.W.L. contributed to preparing the figures; D.W.D., L.C., Y.S., T.W., H.y.C., and W.W.L. contributed to drafting the text.

## Competing Interests

Nothing to report.

## Supplementary Data

**Supplementary Table 1.** Key Resource Table.

| RESOURCE TYPE | RESOURCE NAME | SOURCE | IDENTIFIER | NEW/ REUSE | ADDITIONAL INFORMATION |
| --- | --- | --- | --- | --- | --- |
| Dataset | Simulated mouse primary motor cortex activity during control and decreased pyramidal tract neuron excitability in the parkinsonian condition (RRID:SCR_017571) | DANDI Archive website | https://dandiarchive.org/dandiset/001444 | new | Four complete datasets of simulated mouse primary motor cortex for 6-OHDA experiments. Each dataset includes 4 NWB files (16 files total). |
| Dataset | NeuroMorpho.Org (RRID:SCR_002145) | NeuroMorpho.Org website | https://neuromorpho.org archive name Suter_Shepherd | reuse | Centrally curated inventories of digitally reconstructed neurons and glia associated with peer-reviewed publications |
| Software/code | Python Programming Language version 3.9.18 (RRID:SCR_008394) | Python website | https://www.python.org/downloads/release/python-3918/ | reuse |  |
| Software/code | NEURON version 8.2.2 (RRID:SCR_005393) | NEURON website | https://www.neuron.yale.edu | reuse |  |
| Software/code | NetPyNE version 1.0.5 (RRID:SCR_014758) | NetPyNE website | https://netpyne.org | reuse |  |
| Software/code | Complete code for simulated mouse primary motor cortex 6-OHDA experiments | Github / Zenodo | http://doi.org/10.5281/zenodo.12399983 | new | Detailed primary motor cortex simulation for observing changes in cortical dynamics due to changes in pyramidal tract neuron excitability in parkinsonian mice |

**Supplementary Table 2.** Four datasets, each using different connectivity, stimulation, and neuron location random seeds. Significance (p values) are for Pyramidal Tract (PT5B) neuron firing frequency between conditions (control or parkinsonian) in a particular state (rest or activated).

| Experiment ID | Seeds | Condition | State | Mean PT5B firing rate (Hz) | Standard deviation | Significance (p<value) |
| --- | --- | --- | --- | --- | --- | --- |
| sM1_12-12-2023_01 | conn': 4321, 'stim': 1234, 'loc': 4321 | Control | Rest | 11.7 | 9.4 |  |
| sM1_12-12-2023_02 | conn': 4321, 'stim': 1234, 'loc': 4321 | Parkinsonian | Rest | 14.2 | 7.1 | 0.001 |
| sM1_12-12-2023_03 | conn': 4321, 'stim': 1234, 'loc': 4321 | Control | Activated | 26.8 | 18.2 |  |
| sM1_12-12-2023_04 | conn': 4321, 'stim': 1234, 'loc': 4321 | Parkinsonian | Activated | 29.7 | 12.7 | 0.001 |
| sM1_04-22-2024_01 | conn': 1592, 'stim': 3752, 'loc': 5368 | Control | Rest | 11.3 | 8.0 |  |
| sM1_04-22-2024_02 | conn': 1592, 'stim': 3752, 'loc': 5368 | Parkinsonian | Rest | 19.1 | 9.5 | 0.001 |
| sM1_04-22-2024_03 | conn': 1592, 'stim': 3752, 'loc': 5368 | Control | Activated | 25.7 | 18.4 |  |
| sM1_04-22-2024_04 | conn': 1592, 'stim': 3752, 'loc': 5368 | Parkinsonian | Activated | 29.1 | 11.7 | 0.001 |
| sM1_04-25-2024_01 | conn': 8392, 'stim': 1893, 'loc': 9272 | Control | Rest | 12.4 | 10.2 |  |
| sM1_04-25-2024_02 | conn': 8392, 'stim': 1893, 'loc': 9272 | Parkinsonian | Rest | 22.8 | 11.9 | 0.001 |
| sM1_04-25-2024_03 | conn': 8392, 'stim': 1893, 'loc': 9272 | Control | Activated | 26.4 | 18.3 |  |
| sM1_04-25-2024_04 | conn': 8392, 'stim': 1893, 'loc': 9272 | Parkinsonian | Activated | 30.1 | 12.5 | 0.001 |
| sM1_04-26-2024_01 | conn': 3525, 'stim': 3894, 'loc': 6942 | Control | Rest | 13.4 | 10.3 |  |
| sM1_04-26-2024_02 | conn': 3525, 'stim': 3894, 'loc': 6942 | Parkinsonian | Rest | 24.6 | 12.2 | 0.001 |
| sM1_04-26-2024_03 | conn': 3525, 'stim': 3894, 'loc': 6942 | Control | Activated | 26.6 | 18.3 |  |
| sM1_04-26-2024_04 | conn': 3525, 'stim': 3894, 'loc': 6942 | Parkinsonian | Activated | 30.7 | 12.4 | 0.001 |

**Supplementary Table 3.** Pyramidal Tract (PT5B) neuron parameters in control and 6-OHDA parkinsionian simulations. Ca-V-sensitive big potassium (BK), and transient sodium (NaT) channels changed Current densities in S/cm^2^.

| Condition | BK | NaT |
| --- | --- | --- |
| Control | 7.25x10<sup>-5</sup> | 0.00086 |
| Parkinsonian | 7.25x10<sup>-4</sup> | 0.0172 |

**Supplementary Figure 1.**
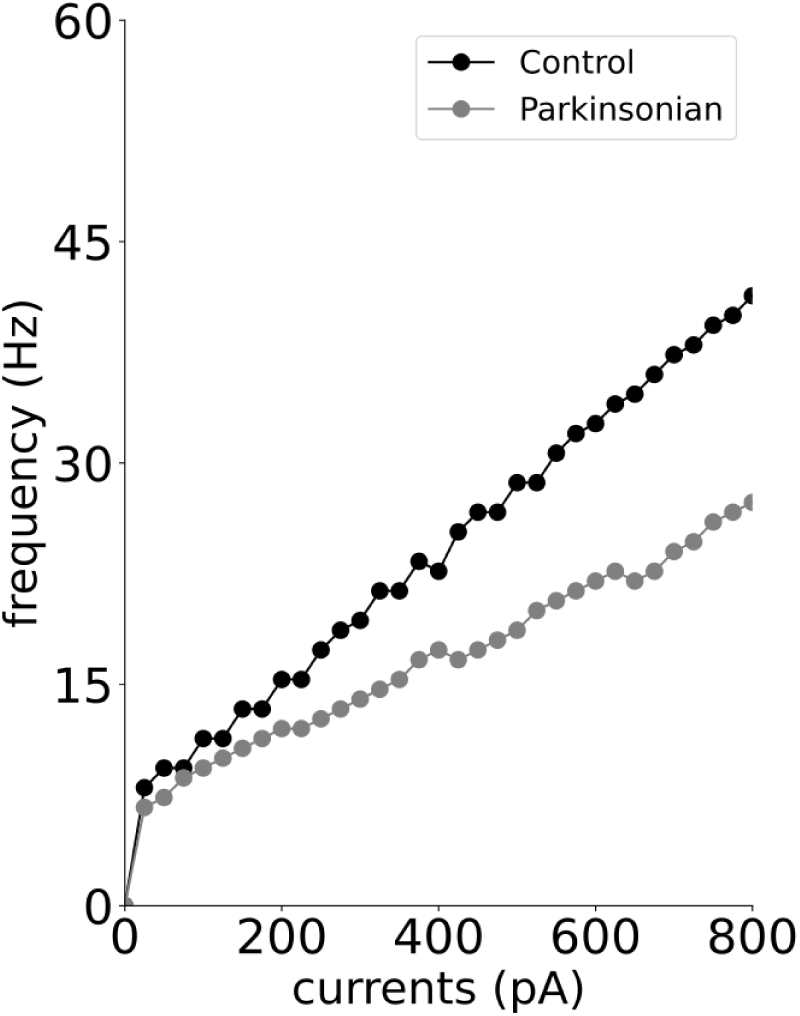
Simulated Pyramidal Tract (PT5B) neuron current frequency curves for control (black) and parkinsonian (gray) conditions.

**Supplementary Figure 2.**
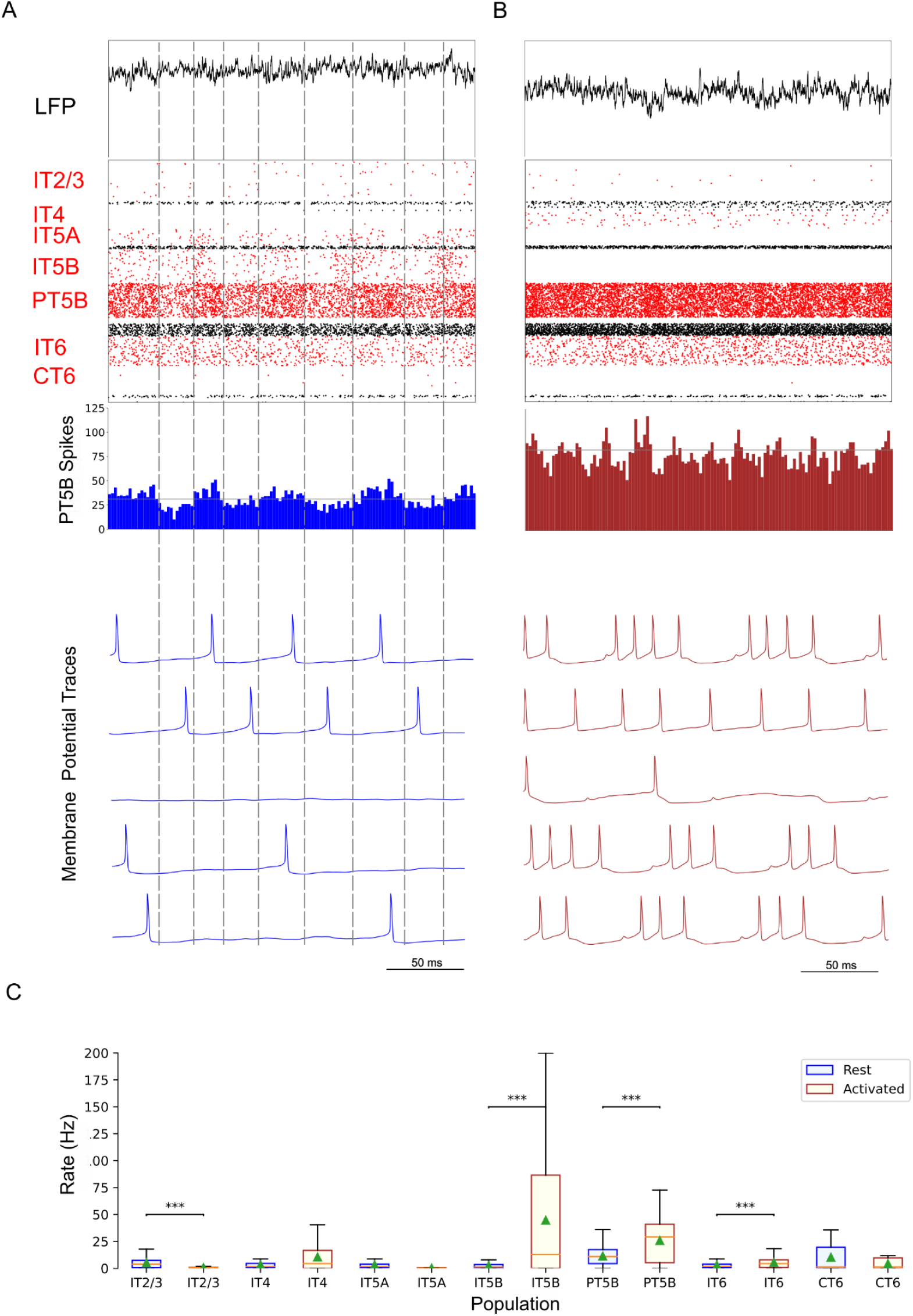
Control condition, simulated primary motor cortex (M1) activity, comparing resting state to movement state. **A.** Resting state (250 ms at midpoint of 4.3 s simulation) displayed dominant beta-band activity (∼20 Hz). Top-to-bottom: local field potential (LFP), raster plot (red excitatory; black inhibitory neurons), PT5B spike count histograms (2 ms bins; dashed lines at half height), and 5 randomly chosen PT5B voltage traces. **B.** Activated state showed change to high-frequency gamma-band activity (∼44 Hz). Same top-to-bottom as in A; bottom traces are the same neurons as in A. **C.** Statistically significant change was found for firing rates for excitatory cell types going from rest to activated: IT2/3 -- rest: 5.0±5.2 spikes/s; activated: 0.64±0.79 spikes/s; PT5B -- 11.3±8.0 vs 25.7±18.3 spikes/s; IT5B -- rest: 2.4±3.0 vs 44.6±59.3 spikes/s; IT6 neurons -- 3.0±3.8 vs 5.3±5.1 spikes/s (mean, std dev, range shown; ***p<0.001; calculated from the last 2.3 s of 4.3 s simulation to avoid initialization transients). sM1_04-22-2024.

**Supplementary Figure 3.**
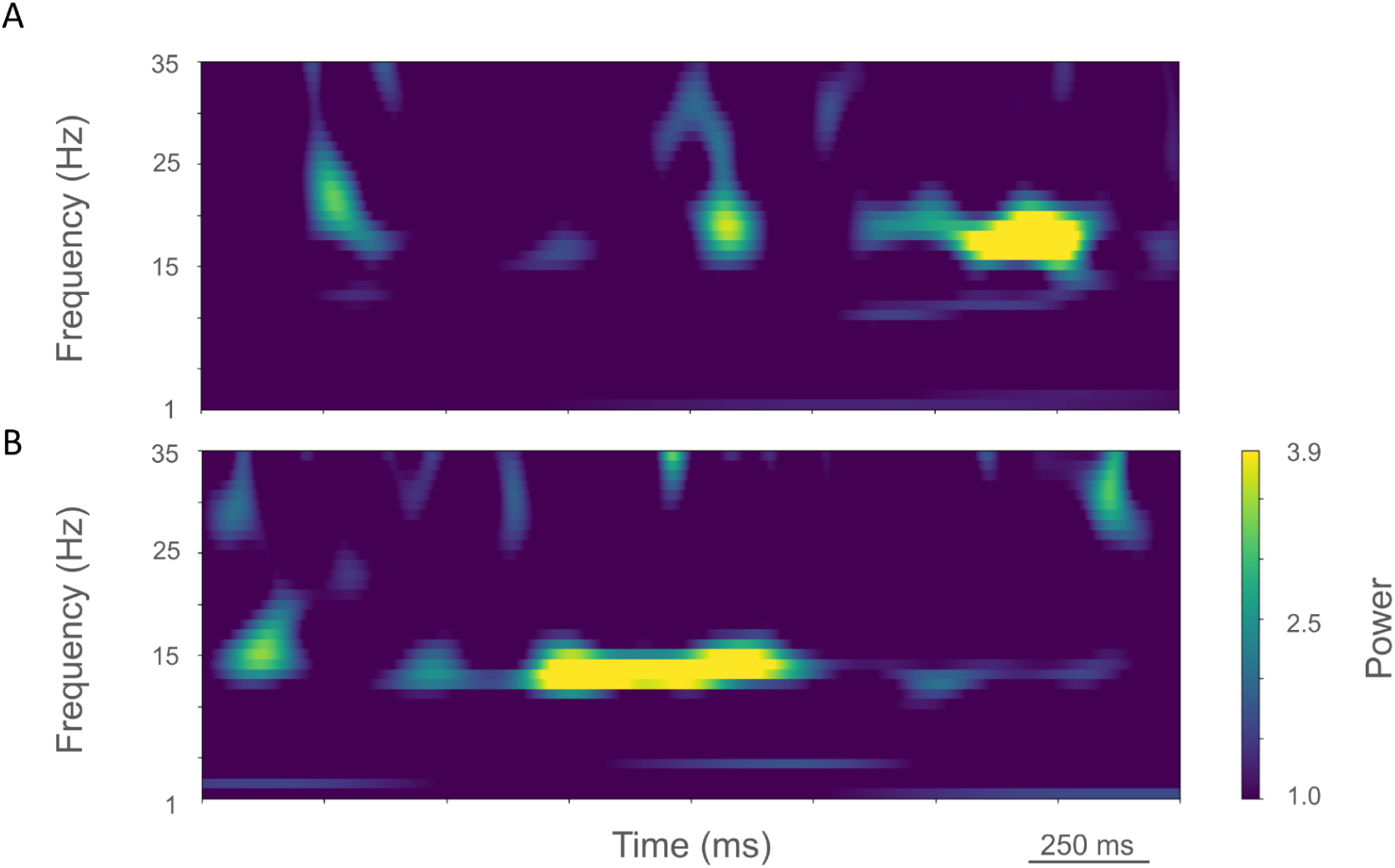
Time-resolved spectrograms of cortical LFPs in M1 in the control condition, under rest and activation states. **A.** The rest state simulation revealed 15-25 Hz beta-band bursts. **B.** During the activated state beta bursts shifted to 15 Hz bursts and higher frequency brief bursts of activity 25-35 Hz are visible. (2.3 s of activity starting 2 s after simulation initiation. Color coding power in bar x 10^−5^). sM1_04-22-2024.

**Supplementary Figure 4.**
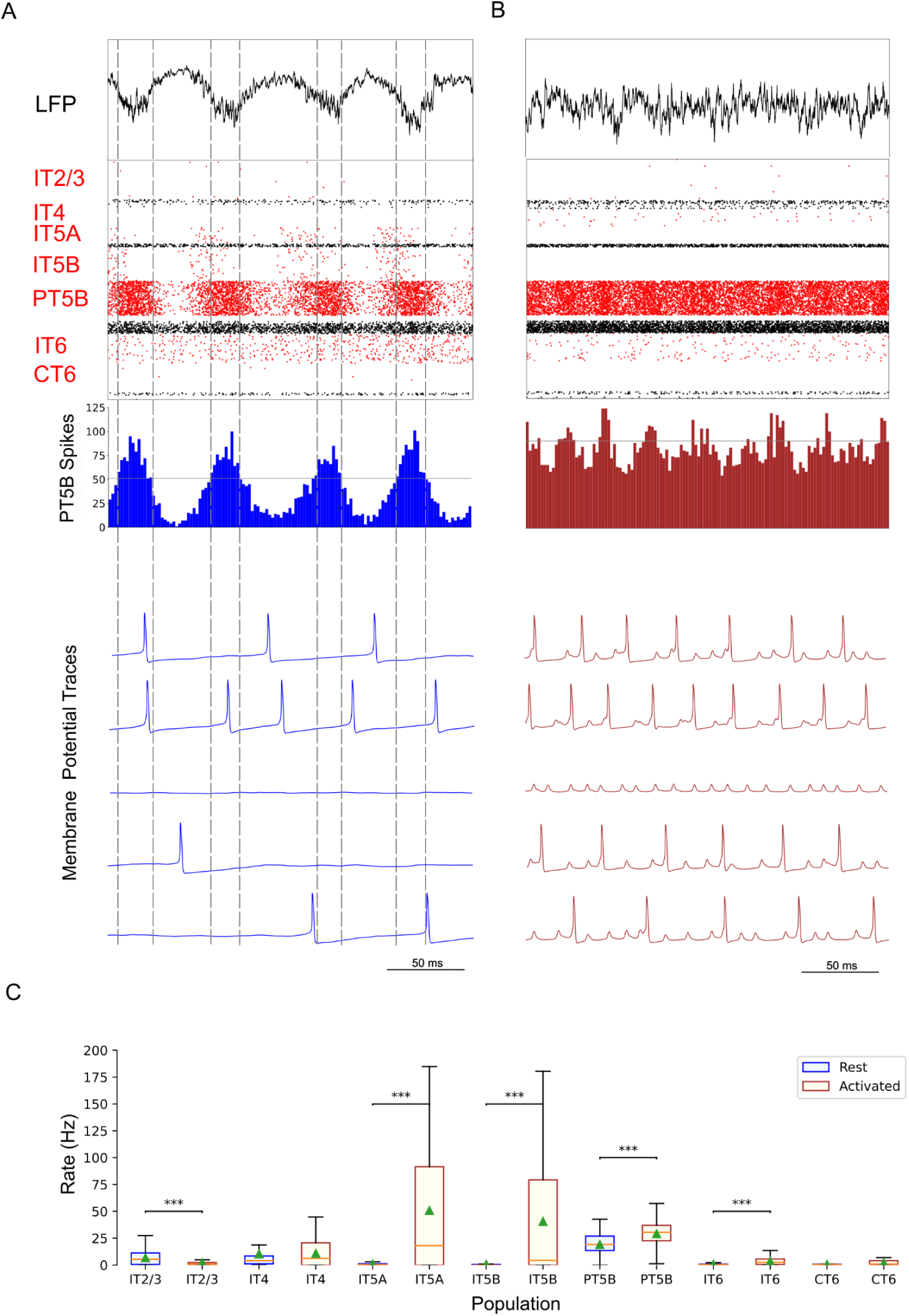
Parkinsonian condition, simulated primary motor cortex (M1) activity, comparing resting state to movement state. **A.** Resting state (250 ms shown at midpoint of 4.3 s simulation) displayed focused 15 Hz beta-band activity. Top-to-bottom: local field potential (LFP), raster plot (red excitatory; black inhibitory neurons), PT5B spike count histograms (2 ms bins; left: dashed lines at half height), and 5 randomly chosen PT5B voltage traces. **B.** Activated state showed change to high-frequency gamma-band activity (∼43 Hz). Same top-to-bottom as in A; bottom traces are the same neurons as in A. **C.** Statistically significant change was found for firing rates for excitatory cell types going from rest to activated: IT2/3 -- rest: 6.8±6.5 spikes/s; activated: 2.1±3.4 spikes/s; IT5A -- rest: 0.89±1.2 vs 50.8±60.5 spikes/s; IT5B -- rest: 0.55±1.2 vs 40.6±55.2 spikes/s; PT5B -- rest: 19.1±9.5 vs 29.1±11.7 spikes/s; IT6 -- rest: 0.76±1.5 vs 3.6±3.8 spikes/s (mean, std dev, range; ***p<0.001; calculated from the last 2.3 s of 4.3 s simulation to avoid initialization transients). sM1_04-22-2024.

**Supplementary Figure 5.**
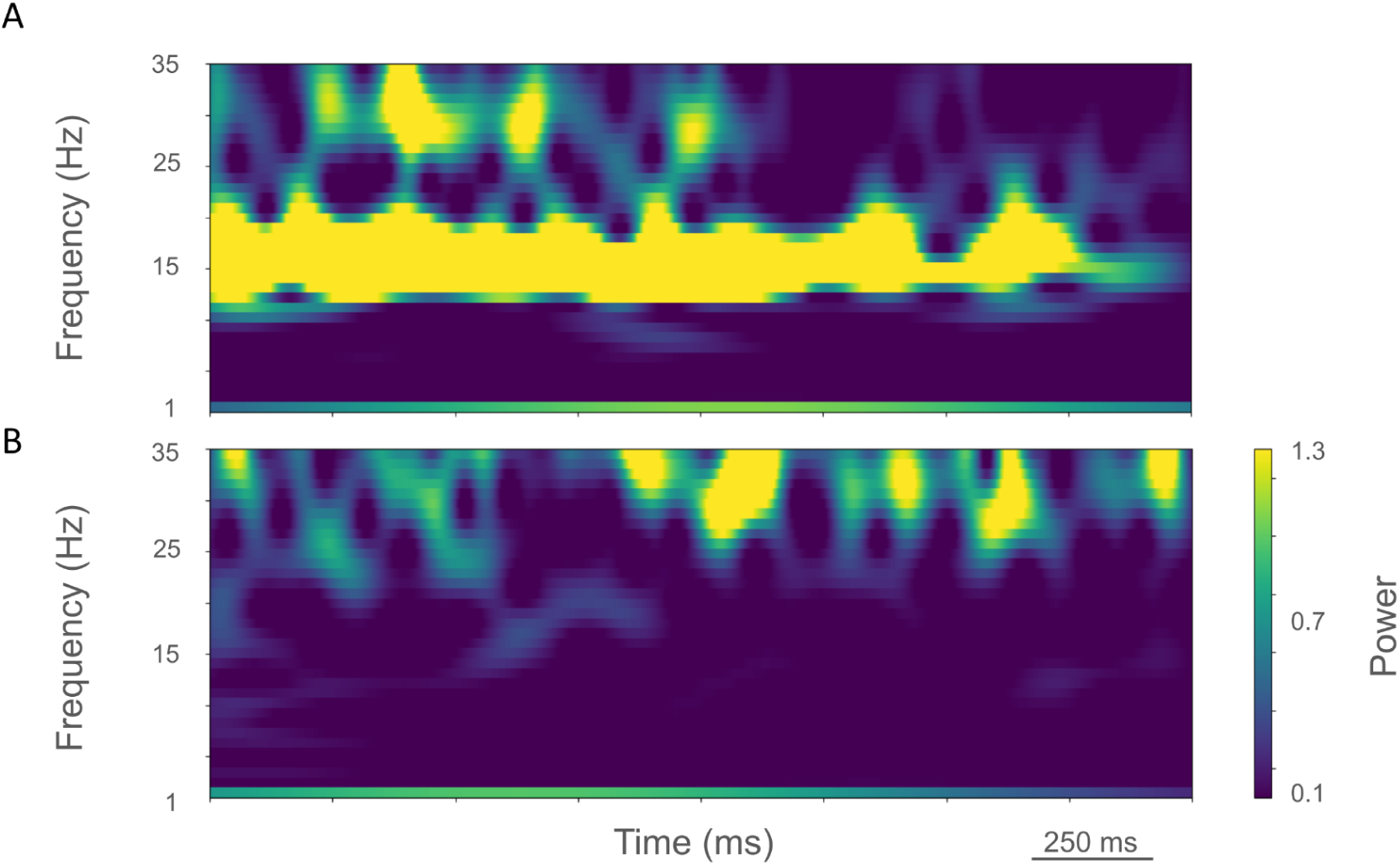
Time-resolved spectrograms of cortical LFPs in M1 in the parkinsonian condition, under rest and activation states. **A.** The rest state simulation revealed a continuous high-power 15 Hz beta-band power with bursts of power in the ∼25-35 Hz. **B.** During the activated state, 15 Hz power was no longer visible and ∼20-35 Hz bursts appeared. (2.3 s of activity starting 2 s after simulation initiation. Color coding power in bar x 10^−4^). sM1_04-22-2024.

## Notes

http://doi.org/10.5281/zenodo.12399983

https://dandiarchive.org/dandiset/001444

https://github.com/suny-downstate-medical-center/pd_m1_6ohda

### Competing Interest Statement

The authors have declared no competing interest.

### Summary of Updates

In this version of the manuscript the description of the model in the Introduction was expanded including how it fits into prior models in the literature. We added substantial detail to the Methods section, especially about connectivity. Additionally, we more fully explain how this model was vetted in our prior publications.

